# Genetically programmable molecular framework for constructing precision recombinant human ECM mimics

**DOI:** 10.64898/2026.10.20.760354

**Authors:** Austin Donnelly Evans, Jenni Keränen, Sasa Rauhalinna, Mandana Mombeinipour, Hasibul Hossain, Paula Turkki, Oommen P. Oommen, Rolle Rahikainen, Vesa P. Hytönen

## Abstract

Engineering extracellular matrix (ECM) mimetic materials that combine molecular precision of synthetic material options with native biological complexity remains a central challenge in biomaterials science. Here, we present a genetically programmable hydrogel platform that enables independent control of mechanical and biological properties including matrix composition, mechanics, viscoelasticity, and biochemical signaling using recombinant human ECM building blocks assembled via isopeptide chemistry. Modular protein ECM crosslinkers containing complete folded ECM domains equipped with terminal SpyTag peptides were integrated within SpyCatcher-functionalized interchangeable natural polysaccharide backbones to generate fully defined matrices with decoupled material parameters. Point mutations on crosslinker SpyTags within the bond interface varied bond affinity from covalent to physical bonding, allowing tuned stress-relaxation behaviors across physiologically relevant regimes without altering matrix composition nor stiffness. Using colorectal cancer models, we show that polymer identity and matrix viscoelasticity independently regulate spheroid growth, invasion capabilities, and dispersion. Importantly, recombinant ECM domains recapitulated native adhesion functions and provide a structurally encoded layer of biological information. This hydrogel platform provides a molecular framework for constructing precision recombinant human ECM mimics with broad applicability across biomaterials science, mechanobiology, disease modeling, tissue engineering, and regenerative medicine.

## Introduction

The extracellular matrix (ECM) is a dynamic and instructive microenvironment that regulates biological processes through coordinated spatiotemporal presentation of mechanical, structural, and biochemical signals.^[1]^ Across developmental states, tissue homeostasis, and diseases, ECM composition and organization continuously gets remodeled by cells, shaping cellular processes including proliferation, migration, differentiation, and survival.^[2]^ Reproducing these complex and evolving microenvironments in vitro remains a central challenge in biomaterials science, tissue engineering, and regenerative medicine.^[3]^ Current approaches to ECM engineering rely predominantly on either synthetic polymer hydrogels or naturally derived matrices from animal or human patient derived sources.^[4]^ Synthetic platforms offer precise control over chemistry, rheological characteristics, mechanics, and material degradation but lack the biological molecular complexity that defines native cellular microenvironments. Hybrid hydrogel systems combine features of both natural and synthetic options, but they typically rely on short peptide sequences rather than full-length domains and native proteins.^[5]^ Peptide sequences like cell adhesive RGD sequences lack the complexity and signaling capacity of fully folded protein domains. Thus, they lose key attributes such as dynamic interactomes and natural mechanical response under cell-borne forces of native ECMs when implemented into a synthetic system.^[6,7]^ Multiple canonical ECM proteins like fibronectin, vitronectin, and fibrinogen contain the same RGD integrin binding sequences.^[8]^ However, differences in ligand structure and folding, integrin selectivity, and receptor organization result in distinct downstream signaling and cellular outcomes despite presentation of the same adhesive motif across different ECM proteins.^[9]^ In contrast, biologically derived matrices and polymers, including collagen, fibrin, tissue-derived decellularized ECMs, and basement membrane extracts such as Matrigel, retain crucial aspects of native tissue structure and bioactivity but suffer from poorly defined composition, batch-to-batch variability, limited tunability, and dependence on animal- or donor-derived sources.^[10,11]^ Consequently, existing hydrogel platforms often impose a fundamental trade-off between engineering control and biological fidelity.^[12]^

Current biomaterial platforms fail to provide molecular building blocks that enable independent and systematic control over individual ECM parameters.^[13]^ Native ECMs comprise of highly diverse networks of proteins, proteoglycans, and glycosaminoglycans whose composition varies between tissues, individuals, and disease states.^[14]^ These components are tightly coupled in conventional natural biomaterials, making it difficult to isolate how specific matrix properties regulate cellular function and find answers to complex ECM biological questions.^[15]^ Parameters typically coupled in existing material systems of broad interest for ECM replication include stiffness, viscoelasticity, remodeling capacity, degradability, and biological cues. As a result, the development of defined human ECM mimetics capable of faithfully and reproducibly recapitulating patient or tissue-specific microenvironments with high biological fidelity has remained elusive.^[16]^ Recombinant protein engineering presents an opportunity to overcome these limitations by treating specific ECM components as genetically programmable material modules. Folded protein domains support biological functions like integrin clustering with synergistic binding sites, cryptic binding sites that become exposed upon mechanical activation, and mechanically activated biological signals. These are connected to architecture of protein domains and co-operation between domains that isolated peptide sequences cannot fully reproduce.^[17]^ Thus, recombinant proteins corresponding to complete protein domains and their combinations would enable precise control over molecular composition, ligand presentation, and intermolecular interactions while eliminating dependence on animal or patient-derived materials.^[18]^ Such systems provide a framework for constructing ECMs from defined molecular components with independently tunable biological and mechanical properties. Furthermore, use of recombinant proteins based on scalable expression systems would offer environmentally friendly and ethical alternatives to traditional methods.

In this study, we introduce a genetically programmable hydrogel platform for the construction of precision-engineered parameter decoupled recombinant human ECM mimetics. The platform integrates a library of human ECM protein domains with selected polysaccharide and glycosaminoglycan backbone components through SpyTag–SpyCatcher-isopeptide chemistry mediated assembly, enabling modular and independent control of matrix architecture, material mechanics, and biological functionality. By varying crosslinker bond interaction strength via single point mutations in SpyTag peptide sequences, switching polymer backbones, and altering ECM crosslinker content within a fully defined molecular framework, we decouple key matrix parameters that are typically inseparable in conventional hydrogels. Using colorectal cancer cells and spheroids as a model biological system, we demonstrate how discrete changes in ECM composition and viscoelasticity differentially regulate tumor cell behavior. We further propose that tertiary folded protein architectures encoded within our protein building blocks constitute an additional and programmable layer of ECM-inspired information, broadening the molecular complexity that can be incorporated into engineered biomaterials. Together, these findings establish a strategy for engineering complex defined human ECMs from genetically encoded recombinant protein building blocks and provide a versatile platform enabling mechanistic studies of cell–matrix interactions across cancer biology, extracellular matrix biology, tissue engineering, and for use in regenerative medicine.

## Results

### Design of programmable recombinant ECM building blocks with decoupled parameters

We created a hydrogel platform that gels spontaneously upon mixing two liquid components in a cell friendly manner. The platform includes both polysaccharide backbone components along with ECM based protein building block crosslinkers. This approach enables the construction of hybrid composite materials with levels of complexity that more closely resemble aspects of the native ECM, while preserving the tunability and modularity of synthetic materials and enabling more independent control of design parameters as illustrated in Figure 1. The first functional parameter of the platform that we wanted to explore included the choice of backbone polymer that could be easily switched without changing other parameters of the system. Dextran and hyaluronic acid (HA) were strategically selected as representative backbone polymers that closely mimic ECM composition while differing in biological function with human cells. As a neutral homopolysaccharide, dextran remains highly bioinert to human cells and does not trigger biological responses. HA, however, is a linear anionic glycosaminoglycan that interacts with multiple cell surface receptors and can be readily degraded by mammalian cells.^[19]^ Both of them are optically clear and thus compatible with several imaging modalities and their biocompatibility is excellent.

**Figure 1:**
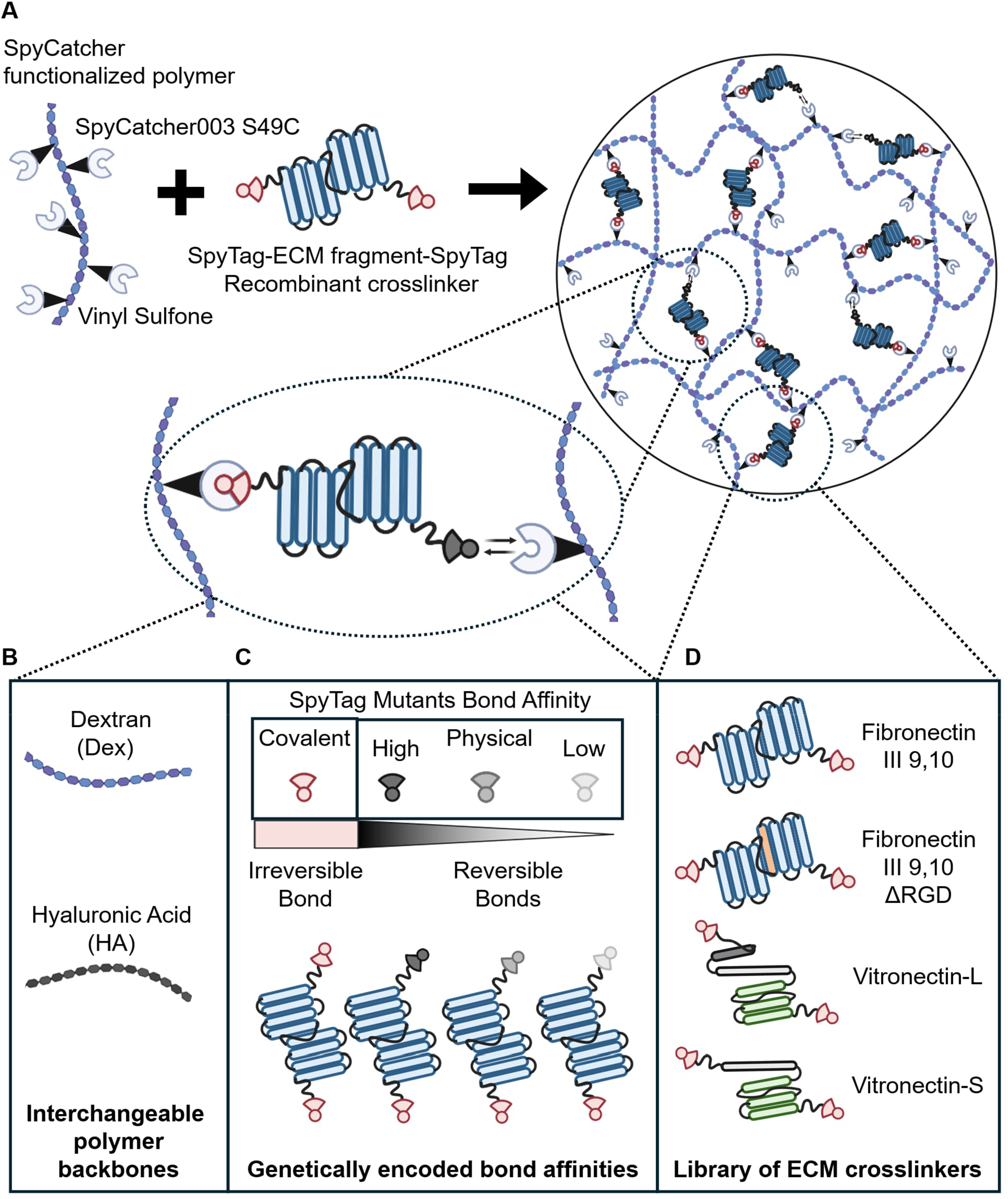
Graphical illustration of hydrogel network design and formation. **A.** Overall architecture of the hydrogel. **B.** SpyCatcher functionalized polymer backbones via vinyl sulfone chemistry. **C.** Library of mutant crosslinkers for controlled viscoelasticity. **D**. Interchangeable ECM crosslinkers to embed biological signals with possibility to include biomechanical characteristics of native proteins.

The second important functional parameter of the material system to program biological functions involved protein engineering ECM adapters for controlled adhesion. We selected well-defined, simple, and robust functional ECM adapters and genetically flanked these segments with tailored SpyTag peptides. We used publicly available protein sequence databases including Uniprot and PDB, to computationally screen multiple domains of human ECM proteins of interest that include different sequences for integrin binding and potential synergy sites. We further experimentally validated that the selected domains can be (1) expressed in *E. coli* for ease of expression scale up, (2) can be purified with affinity chromatography with His-tag NiNTA chemistry, and (3) endotoxins and impurities can be chromatographically removed to avoid any cytotoxic effects towards encapsulated cells. The expression and purification steps are shown with SDS-PAGE in Supplementary Figure S1. We chose human fibronectin III domains 9 and 10 as our starting point due to its known RGD sequence in domain 10, along with synergy site sequence PHSRN in domain 9 for integrin binding and cell adhesion amplification. Further, these domains have known relations and importance to cancer and wound healing.^[20–22]^ To serve as a non-cell binding control with same molecular weight and structure, we also developed the same fibronectin domains with a scrambled AGA in place of the β-integrin-binding RGD sequence (ΔRGD). We further increased our library of ECM crosslinkers with vitronectin 20-292 (Vn-L) and 62-292 (Vn-S). Both adapters contain RGD motifs for cell attachments and hemopexin–like domains (Hp1-Hp3) for heparin, collagen-I and phospholipid binding.^[23]^ In addition, Vn-L contains N-terminal somatomedin B (SMB) domain, a region known to bind PAI-1 and uPAR and contribute to biological processes associated with cell adhesion, migration, and extracellular matrix remodeling.^[24]^

The third important, decoupled parameter that we designed includes the viscoelastic mechanical properties and overall matrix architecture. We leveraged previously reported SpyTag/SpyCatcher mutations that systematically control covalent bond formation and promote controllable non-covalent intermolecular interactions.^[25]^ By integrating these variants into our ECM crosslinkers, viscoelastic properties of the material network could be individually tuned while preserving the overall network architecture, polymer molecular weight, and concentration of all other matrix components. Fusion of the SpyTag variants to recombinant ECM domains generated customizable protein crosslinkers for hydrogel assembly. We positioned a single cysteine residue to position 49 (S49C) to enable attachment of SpyCatcher in highly accessible orientation for isopeptide conjugation via Michael addition to vinyl sulfone click chemistry for bioconjugation.^[26]^

We engineered our hydrogels using 100 kDa dextran and 100 kDa hyaluronic acid functionalized with vinyl sulfone (VS) groups (DexVS and HAVS), providing the carbohydrate backbones for covalent attachment of SpyCatcher003 S49C (SC003-S49C) protein building blocks via thiol-mediated Michael addition to VS moieties. Successful VS conjugation and SpyCatcher coupling was confirmed with NMR (Supplementary Figure S2), and SDS-PAGE (Supplementary Figure S3) A second solution contained the genetically engineered crosslinker ECM building blocks flanked with SpyTag peptides. Recombinantly generated SpyCatcher and SpyTagged ECM crosslinkers were functionally evaluated for their isopeptide conjugation capacity through band shift assay with SDS-PAGE prior to hydrogel formation (Supplementary Figure S4A and B). Crosslinking of the bulk hydrogels occurs spontaneously upon mixing SpyCatcher functionalized polymer solution with SpyTagged ECM crosslinker solution. Viscoelastic features of the generated hydrogels depend on inclusion of mutated SpyTags moieties flanking the ECM crosslinkers. SpyTag/SpyCatcher interactions can span a range of bond stabilities. Wild-type (WT) SpyTag forms an irreversible covalent isopeptide bond with SpyCatcher, generating permanently crosslinked elastic networks with minimal to no stress relaxation. Intermediate viscoelastic behavior denoted as slow relaxing matrices was induced by including non-reactive SpyTag003(D117A; CnaB2 numbering), which abolishes covalent bond formation while retaining high-affinity SpyCatcher binding (K_D_ = 21 ± 4 nM). Importantly, this interaction was previously shown to support talin-mediated force transmission similarly to the covalent SpyTag003/SpyCatcher003 pair, indicating that it remains mechanically stable under physiologically relevant cellular traction forces.^[27]^ The most rapidly relaxing networks denoted as fast relaxing matrices were generated using SpyTag003(D117A/M115A), which exhibits reduced SpyCatcher affinity (K_D_ = 690 ± 84 nM) and forms more transient force-bearing crosslinks.^[25]^ By covalently anchoring one end of every dynamic crosslinker, the hydrogel retains a constant molecular composition over time, decoupling dynamic mechanics from network degradation or crosslinker depletion. Fused to ECM protein domains, these variants enable independent control of matrix composition and network dynamics, spanning irreversible covalent, high-affinity reversible non-covalent, and lower-affinity reversible non-covalent interactions. Together, these mutated isopeptide pairs provide a model system for investigating how molecular bond lifetime and force stability regulate cell-matrix interactions. In the subsequent sections of the paper, we explore the biological outcomes when modulating each individual parameter of the material system while decoupling other material parameters by keeping them constant.

### Genetically programmed building blocks permit custom tissue like viscoelastic material mechanics

Material mechanics were modulated by first altering ECM crosslinker concentration to change the resulting matrix crosslinking density and its storage modulus (G’). Second, material mechanics were altered by including ECM crosslinker -fused mutant SpyTags to influence viscoelasticity and relaxation time while maintaining stiffness across groups as illustrated in Figure 2A. For all gels used in the rheological measurements in Figure 2, polymer concentration was held at 5 wt% DexVS. The concentration ratio of SpyCatcher to SpyTagged ECM crosslinker was maintained at 2:1 as each ECM adapter contains two SpyTag peptides. We conducted oscillation amplitude and frequency sweeps of DexVS matrices with set protein component concentrations to evaluate the linear viscoelastic region and quantify the corresponding storage G’ and loss moduli G”. Concentrations of SpyCatcher:SpyTag ECM crosslinker screened included 1000:500 µM, 1200:600 µM up to 1500:750 µM using SpyTags capable of covalent SpyCatcher binding (Figure 2B). 1000:500 µM ratio generated softer matrices with a storage modulus G’ of 350.0 ± 149.9 Pa, while 1200:600 µM resulted in G’ of 719.3 ± 60.1 Pa, and 1500:750 µM produced stiffer matrices with G’ of 1205.0 ± 127.8 Pa. The loss moduli G” generated from all covalent matrices remained low, and elastic loss tangents < 0.1 were recorded across all groups. This indicates the resulting covalent hydrogels are predominantly elastic with minimal viscous energy dissipation. Table 1 denotes the exact values derived from rheological analyses included in Figure 2B-D.

**Figure 2:**
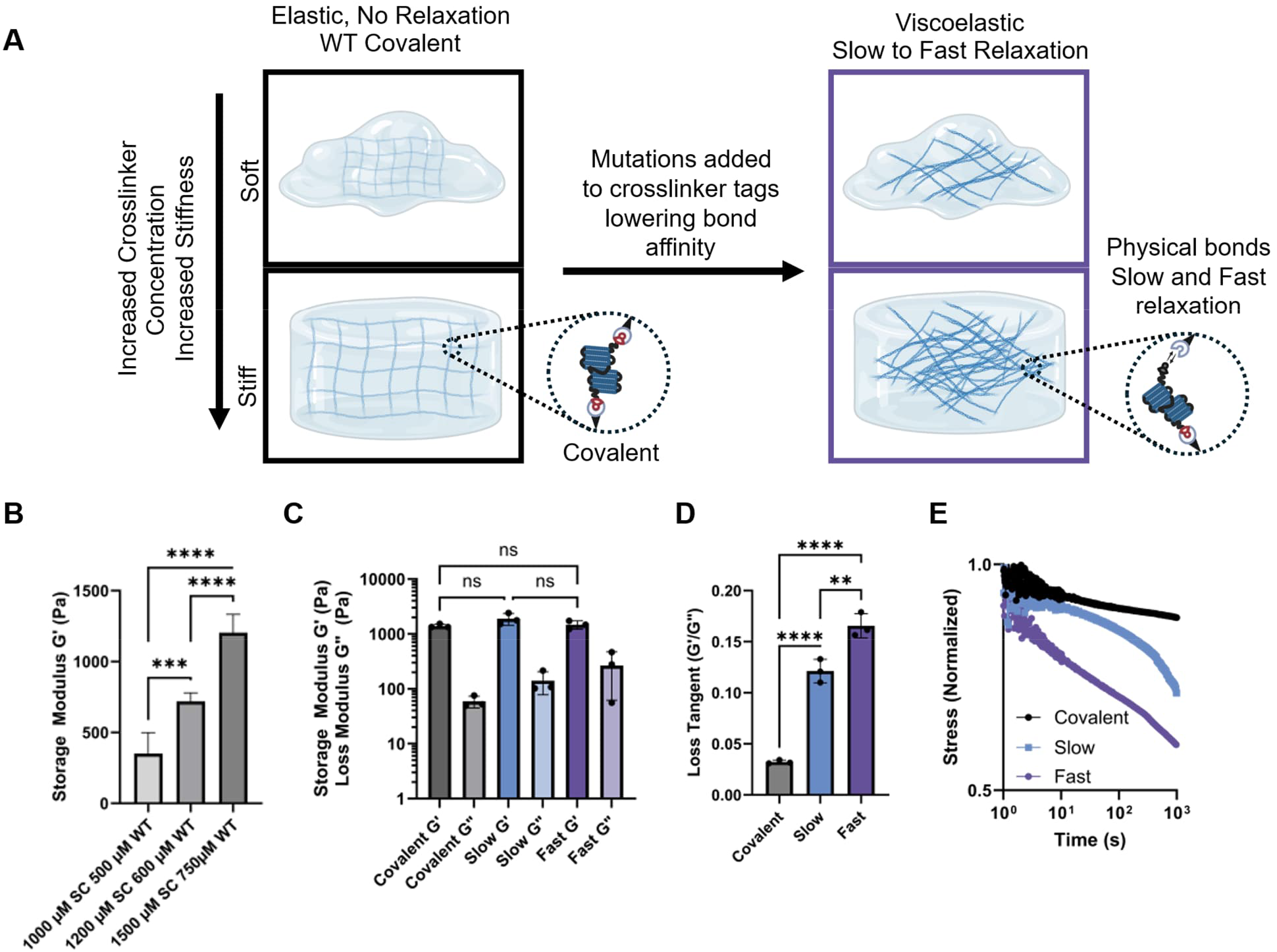
Inclusion of mutant SpyTagged ECM crosslinkers generate modifiable viscoelastic rheological properties of dextran matrices. **A.** Graphical illustration of viscoelastic features and stiffness modulation of hydrogels. **B.** Storage Moduli G’ (Pa) at 1 Hz frequency and 1% oscillation strain of 5 wt% dextran-based matrices with increasing protein crosslinker concentrations, n=3 with mean ± SD illustrated. **C.** Storage moduli G’ (Pa) at 1 Hz frequency and 1% oscillation strain of 5 wt% dextran matrices with mutant spytags, n=3 with mean ± SD illustrated. **D.** Loss tangent (G’/G”) at 1 Hz frequency and 1% oscillation strain of mutant matrices, n=3 with mean ± SD illustrated. **E.** Representative stress relaxation profiles of mutant matrices over time following 15% strain preload, n=3 replicates. For **C-E**, all generated hydrogels were 5wt% DexVS and included 1500 µM SpyCatcher and 750 µM FN III 9,10 crosslinkers with indicated SpyTags (covalent, slow relaxing, and fast relaxing). Data are expressed as mean ± SD. Statistical significance was defined as p < 0.05 and is indicated in the figures by asterisks: * (p < 0.05), ** (p < 0.01), *** (p < 0.001), and **** (p < 0.0001). Group comparisons were assessed using one-way analysis of variance (ANOVA) with Tukey’s multiple-comparison post hoc test.

**Table 1.** Rheological values (mean ± SD) derived from rheological analysis in Figure 2B-E and Figure 4B. 5 wt% DexVS or 2 wt% HAVS backbone with altering ECM crosslinker composition.

| **Group,**<br>**Bond type**<br>**SC:ST Ratio** | **n** | **G' (Pa)** | **G'' (Pa)** | **Loss Tangent,**<br>**Tan $\delta$ ($G''/G'$)** |
| --- | --- | --- | --- | --- |
| Covalent DexVS<br>1000:500 μM | 3 | $350.0 \pm 149.9$ | $21.3 \pm 2.5$ | $0.061 \pm 0.017$ |
| Covalent DexVS<br>1200:600 μM | 3 | $719.3 \pm 60.1$ | $15.0 \pm 0.3$ | $0.021 \pm 0.005$ |
| Covalent DexVS<br>1500:750 μM | 3 | $1205.0 \pm 127.8$<br>(Fig. 2B) | $10.7 \pm 2.2$ | $0.009 \pm 0.017$ |
| Covalent DexVS<br>1500:750 μM | 3 | $1396.0 \pm 114.6$<br>(Fig. 2C-D) | $59.0 \pm 14.3$ | $0.032 \pm 0.002$ |
| Slow DexVS<br>1500:750 μM | 3 | $1899.0 \pm 456.9$ | $141.0 \pm 63.3$ | $0.121 \pm 0.012$ |
| Fast DexVS<br>1500:750 μM | 3 | $1475.0 \pm 251.2$ | $266.8 \pm 205.1$ | $0.166 \pm 0.012$ |
| Covalent HAVS<br>1500:750 μM | 3 | $1040.0 \pm 206.0$<br>(Fig. 4B) | $19.9 \pm 12.0$ | $0.029 \pm 0.010$ |

Next, to generate matrices with altered viscoelastic properties, we used crosslinkers equipped with SpyTags of different binding affinities, while maintaining consistent SpyCatcher and crosslinker concentrations and ratios (1500 µM dextran-immobilized SpyCatcher and 750 µM SpyTag Fn III 9,10 crosslinker; Figure 2C). The resulting storage moduli G’ of all groups remained similar and statistically insignificant between groups with the WT covalent gels having a G’ of 1396.0 ± 114.6 Pa, slow relaxing gels with a G’ of 1899.0 ± 456.9 Pa, and fast relaxing gels with G’ of 1475.0 ± 251.2 Pa. The loss moduli G” and resulting tan delta values, however, significantly changed between groups based on the SpyTag bond affinity incorporated into the matrices (Figure 2D). Covalent matrices exhibited highly elastic behavior with loss tangent (G”/G’) values of 0.032 ± 0.002, which is less than 0.1, the typical loss tangent cutoff for elastic materials.^[28]^ The slow relaxing matrices showed viscoelastic loss tangent values of 0.121 ± 0.012, and the fast-relaxing matrices displayed higher loss tangent values of 0.166 ± 0.012 both resembling typical soft biological tissues such as liver and spleen tissues.^[29]^ The stress relaxation profiles in Figure 2E revealed pronounced differences in matrix viscoelasticity. Covalent gels behaved predominantly elastically and maintained stress over timescales exceeding hours (10³ s). By comparison, slow-relaxing matrices exhibited estimated stress relaxation half-times on the order of hours, while fast-relaxing matrices dissipated stress within minutes, corresponding to the 10² s timescale.

### Interchangeable ECM crosslinkers drive cell adhesion and spreading differently

Cell interaction with ECM domains through integrin binding enables directed and controlled cell adhesion profiles to the matrices. Figure 3A illustrates the design considerations behind the selection of each adapter employed in the first library of ECM mimetic crosslinkers from full length fibronectin and vitronectin. We employed 2D cell culture adhesion assays with mouse embryonic fibroblasts (MEFs) to test cell adhesion profiles of each individual crosslinker and compare their ability to support cell adhesion with their native full length protein counterparts. CyQuant (Figure 3B) and crystal violet (Supplementary Figure S5A) were used to quantitate the number of cells attached to each biofunctionalized substrate. Bovine Serum Albumin (BSA) coating was employed as a negative control not supporting the binding of fibroblast cells. FnIII9,10 adapters exhibited adhesion profile comparable to full length fibronectin protein with insignificant differences. When the RGD sequence in FnIII10 domain was mutated to AGA, cell adhesion was markedly reduced, reaching baseline levels observed with BSA, a non-adhesive control that lacks integrin-binding capability. Vitronectin adapters L and S revealed similar adhesion profiles trending towards slightly reduced adhesion profiles compared to full length vitronectin and to full length fibronectin. Figure 3C shows representative brightfield images of MEF cell adhesion profiles as quantitated in Figure 3B including cell density on the substrate and zoom in revealing adherent cell morphology with each crosslinker coating the substrate.

**Figure 3:**
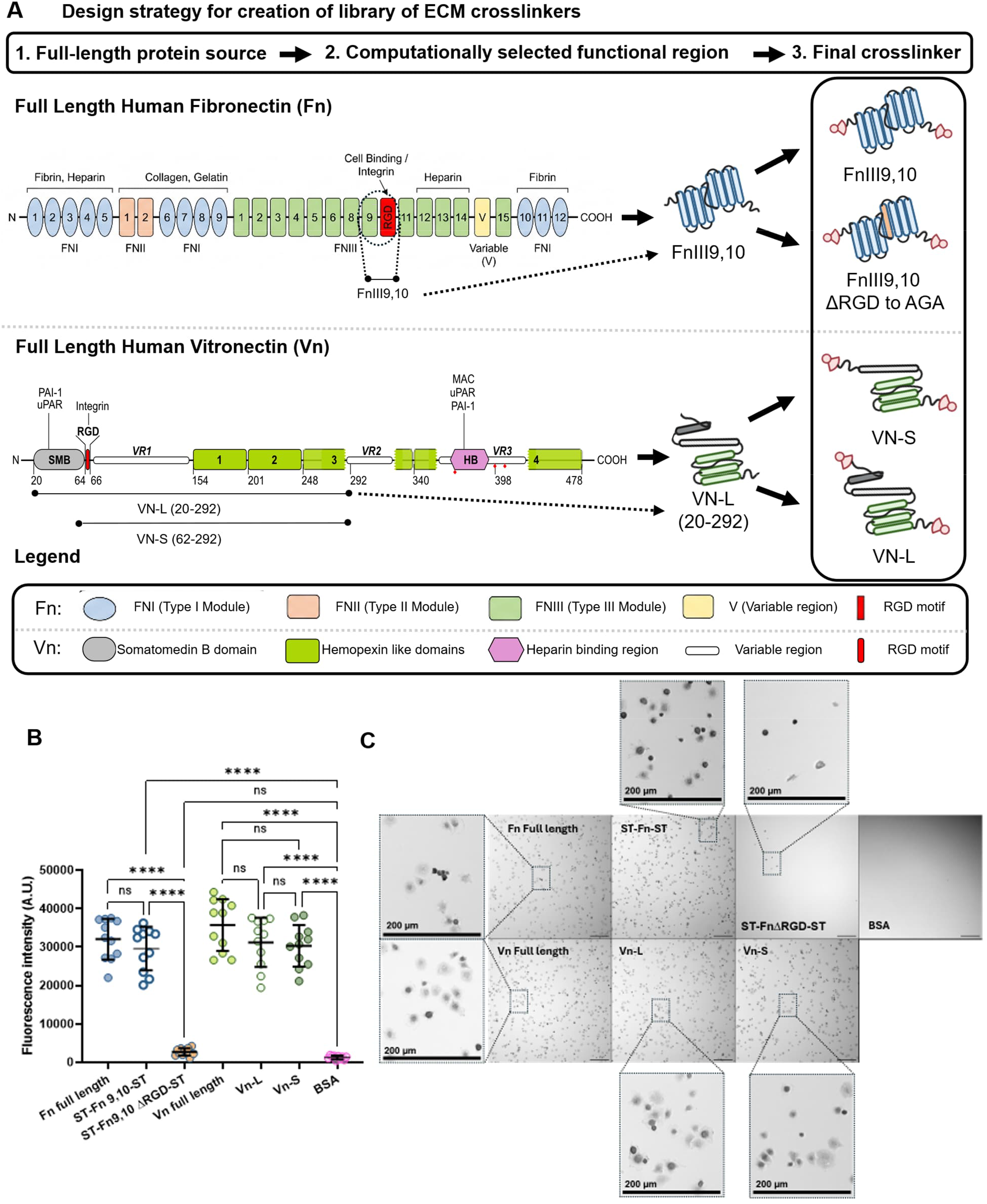
Design and adhesion characterization of recombinant ECM crosslinkers. **A.** Graphical illustration of initial ECM crosslinker library and design considerations from full length human fibronectin (adapted from^[30]^) and human vitronectin (adapted from ^[31]^). **B.** MEF cell adhesion profile based on fluorescence measured with CyQUANT assay, n = 3 biological replicates. Data are expressed as mean ± SD. Statistical significance was defined as p < 0.05 and is indicated in the figure by asterisks: * (p < 0.05), ** (p < 0.01), *** (p < 0.001), and **** (p < 0.0001). Data distribution normality was evaluated using the Shapiro-Wilk test. For normally distributed CyQUANT datasets (n = 3), a one-way Analysis of Variance (ANOVA) followed by Bonferroni’s post-hoc test was performed. **C.** Representative brightfield images (Olympus IX51) showing MEF attachment, inserts with zoom in on morphology, and cell density across substrate conditions following 20 min incubation (scale bars = 200 µm).

### Hyaluronic acid pushes colorectal cancer cells towards invasive phenotypes enabled by polymer backbone switching

We employed colorectal cancer cell lines as a model cell system to delineate the effect of modulating individual material parameters in a simple cancer invasion assay setup. We switched the polymer backbone of the material system illustrated in Figure 4A without significantly altering material mechanics within a fully defined covalently crosslinked and elastic system (Figure 4B). Storage moduli of generated Dex and HA based matrices remained similar at 1088.0 ± 5.0 Pa and 1039.0 ± 206.0 Pa respectively. HCT116 encapsulated in bulk dextran and HA hydrogels exhibited completely different visual phenotypes after 7 days of culture with high proliferation and viability of >90% (Figure 4C). In dextran-based matrices, cells retained spheroidal shape and integrity throughout the culture period, while in HA matrices, the HCT116 cells separated into single cells and migrated away from each other, exhibiting disperse phenotype.

**Figure 4:**
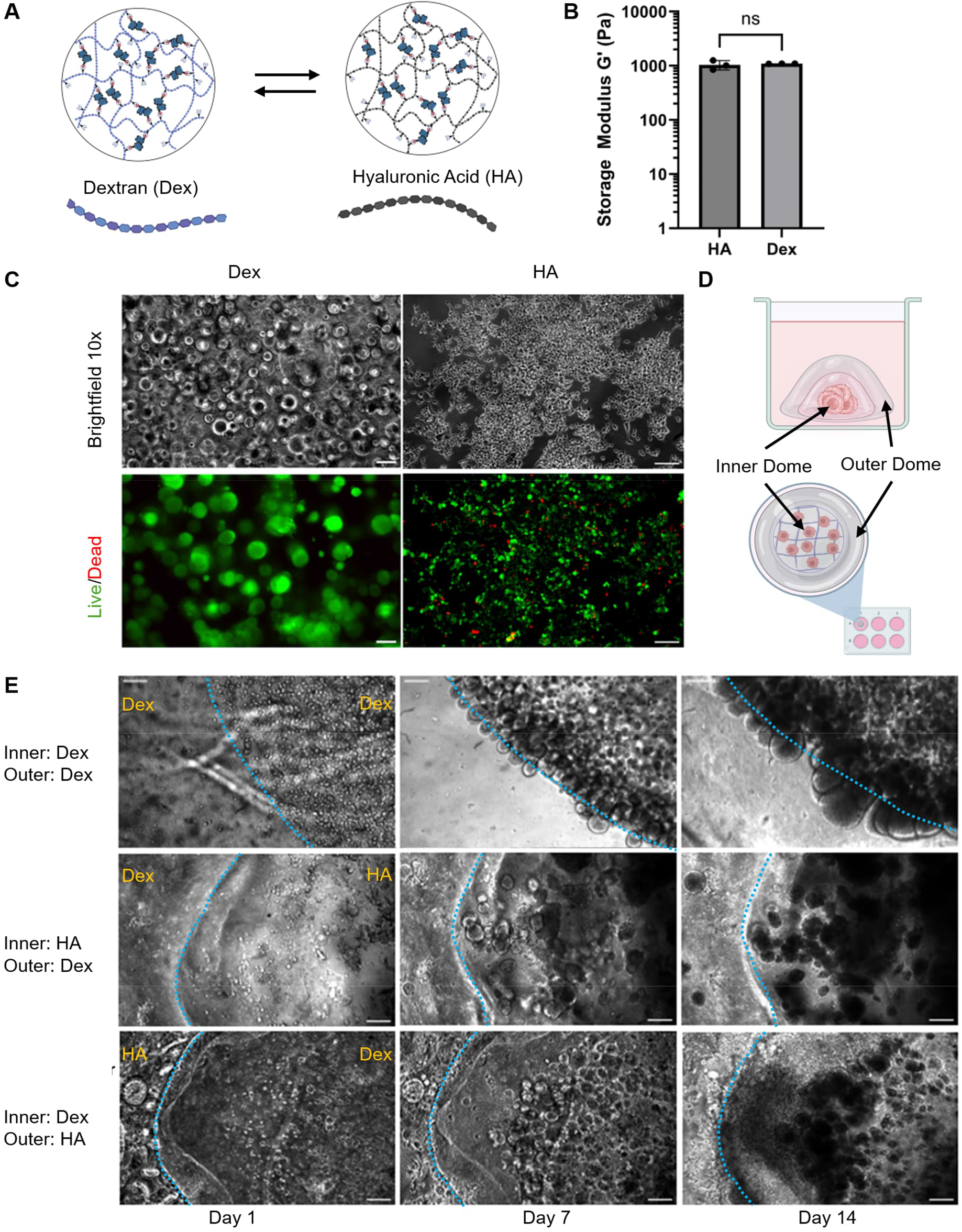
Backbone polymer identity dictates HCT116 colorectal cancer spheroidal integrity and invasion capacity. **A.** Graphical illustration of backbone polymer change. **B.** Storage moduli G’ (Pa) of 5 wt% DexVS and 2 wt% HAVS matrices at 1 Hz frequency and 1% oscillation strain. Data is expressed as mean ± SD. Unpaired t-test was performed to evaluate variance between groups. **C.** Representative brightfield and live dead microscopic images of HCT116 cells encapsulated in Dex and HA matrices at day 7 taken at 10x, scale bar = 200 µm. **D.** Graphical illustration of dome invasion assay setup. **E.** Representative brightfield images over time from day 1, day 7, and day 14 of HCT116 cells encapsulated in domes of Dextran and Hyaluronic Acid hydrogels at 10x.

To further assess the biological effects of the polymer backbone, we next employed a simple hydrogel “dome-within-a-dome” setup with HCT116 colorectal cancer cells encapsulated in the middle dome at the start of the experiment illustrated by Figure 4D. We switched the base polymer of each dome in different orders and analyzed how the cells invaded and behaved through the created material boundaries over the course of 14-day culture periods. When HCT116 were encapsulated in fully dextran-based hydrogels, the HCT116 cells were unable to spread and break through the material barrier within 14 days, growing in spheroid-like structures at the material barrier. When plated within an HA based matrix and then further surrounded by dextran-based gel, the cells formed spheroids within the HA matrix, but did not penetrate through the dextran, nor did they reach the barrier like they did in fully dextran matrices. When the cells were plated in dextran and surrounded by HA gel (Figure 4E), they formed spheroid structures within the dextran base material, but invaded out through the HA and spread into single cell like phenotypes as seen above in the HA alone matrix (Figure 4C).

### Viscoelastic control of matrices elucidates spheroidal formation and invasive properties of colorectal cancer

We next evaluated the effect of viscoelasticity alone as a biological material parameter on HCT116 spheroid growth. By modulating the viscoelastic properties of the hydrogel system with altering bond strengths from covalent to slow relaxing and fast relaxing matrices (Figure 5A), we elucidated morphological differences in spheroid formation and growth as shown by representative images in Figure 5B. HCT116 cells seeded as single cells grew into spheroids over the course of 7 days in culture. From the formed spheroids, we then utilized image-based analyses to quantify structural properties like spheroid size (Figure 5C) and circularity (Figure 5D). HCT116 cells in covalent elastic matrices with and without RGD present exhibited confined, dense spheroids that remained smaller in size in the culture time frame. In slow relaxing matrixes, in addition to the confined spheroids, we observed some slightly larger and more irregularly shaped spheroids. This phenomenon was most noticeable in the fast relaxing matrices, where more of the spheroids grew significantly larger with noticeably irregular rough forms, when compared to the WT covalent elastic matrixes. While the circularity of the main spheroid population remained consistent across all groups, more irregular morphologies were observed when matrix stress-relaxation rate increased.

**Figure 5:**
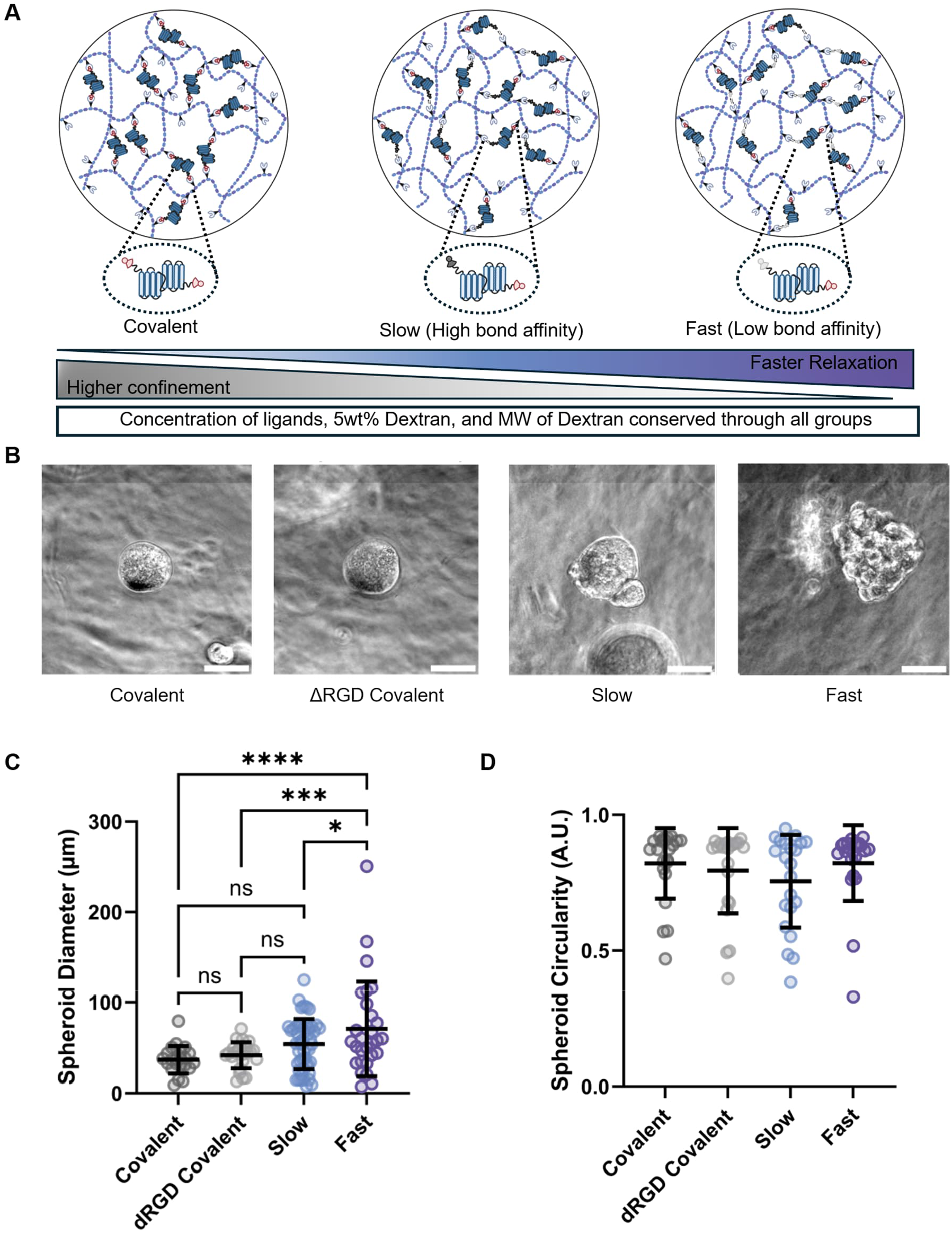
Viscoelasticity of matrix influences colorectal cancer spheroid size and shape. **A.** Graphical illustration of modulating viscoelasticity by changing crosslinker bond type. **B.** Representative brightfield images of encapsulated HCT116 spheroids on day 7 taken at 20x magnification. Scale bar = 50 µm. **C.** Quantified spheroid diameter (µm) of HCT116 colorectal cancer spheroids at day 7, n = 22. **D.** Spheroid circularity of HCT116 colorectal cancer cells measured at day 7, n=22. For C and D, data is expressed as mean ± SD. Shapiro wilks normality followed by One-Way ANOVA analysis of variance tests were conducted. Statistical significance was defined as p < 0.05 and is indicated in the figure by asterisks: * (p < 0.05), ** (p < 0.01), *** (p < 0.001), and **** (p < 0.0001). For **B-D**, all generated hydrogels contained 5 wt% DexVS with SpyCatcher to respective group SpyTag Fn III 9,10 crosslinker ratio of 1250:625 µM with 0.5 mg/mL supplemented collagen type I to help hold matrix integrity through experiment timelines.

## Discussion

This study establishes a genetically encoded recombinant material platform that enables systematic decoupling of matrix physiochemical properties individually from one another. Such modularity addresses a longstanding challenge in biomaterials design, where modifying one matrix parameter often unintentionally alters others, especially in biologically relevant parameters. First, independent control over material mechanical properties like stiffness and matrix viscoelasticity is particularly important because these properties directly influence cellular behaviors like cell spreading, migration, differentiation, and matrix remodeling.^[32]^ Our results show that the mechanical properties of this framework can be modulated across biologically relevant ranges by altering the protein concentrations and altering the bond strength with selected point mutations in the SpyTag-SpyCatcher binding region. Mechanical tuning can further be achieved without altering ECM crosslinker type nor identity. By allowing mechanical tuning while maintaining comparable ECM compositions, this platform provides a useful tool for isolating mechanical dependencies in biological systems. Furthermore, the ability to exchange polymer backbones while preserving overall hydrogel mechanical properties and architecture highlights added versatility to this material system design. This feature permits material customization to specific biological or translational requirements without necessitating complete redesign of the recombinant ECM crosslinker components themselves.

Interchangeable ECM crosslinkers provide another design advantage and biochemical information dimension beyond bulk material mechanical control. Because protein properties and folding can influence signaling, adhesion site presentation, and mechanical accessibility, this platform offers opportunities to investigate how cells respond not only to ECM composition but also to specific folded protein domains, mechanobiological sensitization, and added informational layers outside of the possibilities offered by peptide sequences alone.^[33]^ Different ECM domains may influence cellular adhesion profiles, downstream signaling cascades, and cellular remodeling in ways that current peptide based or natural materials cannot provide enough decoupling to investigate methodically.^[34]^ Additionally, the modularity of the system enables proteins to be incorporated through one or both termini, potentially allowing the tuning of force transmission pathways and the mechanical loading experienced by individual domains. Such designs could facilitate the incorporation of mechanically responsive proteins containing cryptic binding or signaling sites, anchor morphogens and growth factors to simulate ECM sequestration, or provide opportunities to examine how force-dependent molecular unfolding regulates cellular behavior.^[35]^ Moreover, the platform may enable the replacement of naturally weak or transient molecular interactions with engineered covalent binding pairs, offering a means to systematically study the contribution of interaction stability to matrix remodeling and mechanotransduction.^[36,37]^ Looking forward, the same chemistries could potentially be adapted to interface directly with cells through genetically encoded or chemically introduced binding tags generated from inside a genetically reprogrammed cell, creating opportunities to investigate cell-mediated matrix organization, focal adhesion recruitment processes, and force transmission with unprecedented control from both sides of the cell membrane.^[38]^ Nevertheless, the recombinant matrices generated herein possess a fully defined composition and lack the exogenous growth factors and poorly defined constituents present in commonly used materials such as Matrigel, offering a more controlled microenvironment for disease modeling and drug screening applications.^[39]^

Moreover, the ability to control and modulate viscoelasticity of the generated matrices precisely by altering the bond strength of bio-orthogonal isopeptide forming proteins with genetically encoded point mutations further increases the power and biological relevance of the platform. Native tissues are viscoelastic in nature, and viscoelastic matrices with controlled stress relaxation profiles mimic these environments more closely than existing fully covalent elastic hydrogels.^[40]^ Viscoelastic features further affect cellular mechanotransduction pathways through stress relaxation and time dependent force dissipations on cellular levels.^[41,42]^ The stress relaxation profiles exhibited by the matrices generated in this study further align with biologically relevant viscoelastic dynamics. By comparison, fast-relaxing matrices dissipated stress on the order of minutes (10² s), a timescale associated with processes such as focal adhesion turnover, cytoskeletal reorganization, and short-term force adaptation^[43]^, whereas slow-relaxing matrices retained stress for hours, approaching timescales relevant to matrix remodeling, cell spreading, migration, differentiation, and longer-term tissue organization.^[44]^ This platform enables altering the viscoelasticity with genetic mutations rather than bulk material properties like molecular weight of polymer, changing of chemical crosslinking bond types, and concentrations or densities of components within a common framework. This facilitates more rigorous investigation of cell-matrix interactions and improves experimental reproducibility across biological and mechanical parameters. The modular nature of this platform makes it particularly suitable for future studies requiring systematic investigations of matrix cues like stem cell fate regulation, disease modeling with the possibility to explore specific protein mutations or damage in the ECM, organoid culturing, and regenerative medicine applications without the need for natural sources like patient or animal derived materials.

Several limitations resulting from this work and the platform should be acknowledged. While multiple matrix parameters were independently varied in this study, only selected polymers and a small number of ECM regions were evaluated with a single representative cancer cell type to illustrate the power of genetically programmed decoupled parameters of the material system. Moreover, expressing complex and large ECM proteins requiring extensive post translational modifications like collagens and laminins remains challenging.^[45,46]^ Future work should explore protein production in different expression systems to enable more complex crosslinker designs with post translational modifications involved. Likewise, further improvements and scale up in recombinant protein production could support the development of larger biofabricated tissue constructs and organ-scale systems, though achieving sufficient protein production scale remains a challenge in current academic settings. Future studies should investigate broader ECM domains and crosslinker protein libraries with large scale recombinant production, long term remodeling behaviors of the materials, cellular phenotypic behavior with transcriptomics, and future potential in vivo applications.

In conclusion, this genetically programmable hydrogel platform enables independent tuning of polymer chemistry, mechanical properties, ECM biofunctionality, and viscoelastic behavior. This design framework addresses the key challenge of decoupling matrix variables that are traditionally intrinsically linked together while incorporating human ECM protein complexity and additional informational layers beyond what is possible with peptide sequences alone. This platform provides a powerful structure for the creation of higher order complexity human mimetics and generating vast arrays of different types of materials without the need for animal or human patients derived materials.

## Materials and Methods

### Materials

100 kDa Dextran was procured from Sigma Aldrich. 100 kDa Hyaluronic Acid was procured from Lifecore biomedical, Chaska Minnesota USA. All buffers, acids, bases, chemicals, and solvents used were of analytical quality sourced from Sigma Aldrich.

### Plasmids and Cloning

The genes of interest were inserted into expression plasmid pET-28(+)a containing T7 promoter for target protein expression and kanamycin resistance gene for selection. Target ECM protein’s sequences were retrieved from UniProt or NCBI database and codon optimized for expression in *E. coli* utilizing GeneArt GeneOptimizer service by Thermo Fisher Scientific. Then the genes were synthesized and cloned into the pET28a(+) expression plasmid by GenScript service. The expression plasmid for SpyCatcher003(S49C) was a kind gift from Prof. Mark Howarth (University of Cambridge) and has been previously described.^[26]^ Protein expression constructs will be deposited into Addgene and released for open source scientific research use.

### Protein Expression

Expression plasmids were transformed into chemically competent *E. coli* BL21 star DE3 cells by heatshock at 42 °C for 30 seconds. Transformed bacteria were grown on lysogeny broth (LB) agar plate with kanamycin overnight at 37 °C. Seed cultures were prepared by inoculating a single colony to seed culture media (LB media with 0.8 % (v/v) glucose and 50 µg/ml kanamycin) and incubating them overnight with 225 rpm agitation at 37 °C. 1 L expression culture was done in 2.8 L flasks with the same media as in seed culture. The bacteria were grown up to 0.8 OD_600_ at 37 °C with 225 rpm agitation. The protein expression was induced using 0.2 mM isopropyl β-D-1-thiogalactopyranoside (IPTG). After the addition of IPTG, the temperature was lowered to 18 °C and allowed protein expression overnight. After protein expression, bacterial cells were harvested by centrifuging at 5000 g for 30 minutes at 4 °C. To evaluate feasibility of large-scale manufacturing, *E. coli* expression was successfully scaled up from 1 L flask culture to 5 L Labfors Infors bioreactor using defined media and pO-controlled fed-batch process.

### Protein Purification

Harvested bacteria were resuspended in binding buffer (50 mM Tris, 500 mM NaCl, 20 mM imidazole) and 1 tablet Pierce Protease inhibitor (Thermo Fisher) per 0.25 L buffer, 50 µg/ml lysozyme, and 2 mM MgCl_2_ were added. Proteins were released from the cells by sonication at 4°C for 5 minutes with 2/5 second on/off cycles at 60% amplitude. Soluble ECM proteins (Fn and Vn), were separated from cell debris and insoluble proteins by centrifugation at 20,000 g for 15 minutes at 4°C. Cleared lysates were then incubated with complete His-tag Purification Resin (Roche) on roller shaker for 1 hour at 4°C. Purification continued on BioRad’s open column, where unbound protein was removed and the resin was washed with 15 ml binding buffer (50 mM Tris-HCl, pH 8, 500 mM NaCl, 20 mM Imidazole) followed by flush with endotoxin releasing wash buffer (50 mM Tris-HCl, pH 7.5, 150 mM NaCl, 0.1% Triton-X114). Finally, target proteins were eluted by elution buffer (50 mM Tris pH 8, 500 mM NaCl, 250 mM imidazole). Eluted proteins were subsequently dialyzed to 1x TBS (50 mm Tris-HCl, 150 mm NaCl, pH 8) buffer at 4°C, aliquoted, concentrated with 10 kDa MWCO centrifugal filters (Amicon Ultra, Merck) and stored at -80°C.

### Vinyl Sulfonation Chemistry onto Polymer Backbone

100 kDa dextran or 100 kDa HA were dissolved in 0.1M NaOH in diH_2_O at 2% w/v into 100mL (2g dextran or HA in 100 mL at 20 mg/mL concentration) under constant stirring. Divinyl sulfone was added dropwise at a 1.25 molar excess relative to dextran glucose monomer units (1.55 mL of divinyl sulfone added for dextran, 625 µL of divinyl sulfone added for HA reaction). Reaction was allowed to stir for 5 min and then quenched by adding 10M HCl dropwise and adjusting pH to below 5. After reaction, the reaction mixture was dialyzed against diH_2_O for 3 days with 3x water changes per day. After 3 days dialysis, DexVS solution was pH adjusted to neutral pH 7. DexVS was dialyzed for a further 24 hours after pH adjustment, and solutions were frozen and lyophilized. Degree of VS modification was analyzed and confirmed using ^1^H-NMR in deuterated H_2_O. ^1^H-NMR spectrum recorded at 298 K using a JEOL NMR spectrometer.

### Gel Formation

Lyophilized DexVS and HAVS were weighed and dissolved in PBS at batch-specific volumes calculated to achieve desired final polymer and protein concentrations. SpyCatcher003 S49C was added at a volume determined from measured protein stock concentration to dissolved polymer solution and allowed to sit for at least 30 minutes at room temperature to allow Michael addition coupling. Hydrogel precursor solutions were prepared using volumes calculated individually for each batch to maintain constant final polymer, ligand, and crosslinker concentrations despite variations in stock concentrations. A concentration calculator was used for all formulations. SpyCatcher-polymer precursor solution was pipetted into mold or free droplets into well plates. ECM crosslinker solution was pipetted into polymer droplet and mixed with gentle pipetting up and down along with tip mixing simultaneously to initiate crosslinking. For generating cell laden matrices, counted cell pellets were suspended and mixed with SpyCatcher-polymer precursor solutions prior to adding ECM crosslinker protein solutions.

### Rheology

200 µL gels were formed in 10 mL, 12 mm diameter syringe molds and allowed to crosslink overnight prior to rheological measurements. All rheological evaluations were conducted on a TRIOS Discovery HR 2 rheometer at 37°C using 12 mm parallel plate geometry with solvent trap to maintain sample hydration throughout measurement periods. The general rheological protocol followed for all fully crosslinked gels started with 0.5 N axial force pre-straining between plates after sample loading followed by amplitude sweeps from 0 to 20% oscillation strain at fixed 1 Hz frequency followed by frequency sweeps from 0 to 10 Hz at fixed 1% oscillation strain. Loss tangent or tan delta was derived from storage and loss modulus values from frequency sweeps at 1 Hz and 1% oscillation strain. Stress relaxation analysis was then conducted for up to 12 hours with 15% applied initial strain and measuring resulting modulus over time, G(t).

### Cell Culture

HCT116 cells acquired from ATCC were cultured following normal tissue culture protocols in T75 flasks with complete DMEM media (1x DMEM Glutamax (Gibco), 10% fetal bovine serum (Gibco), 1% penicillin/streptomycin as added antiobiotic (Gibco)). Cells were cultured in cell incubator at 37°C and 5% CO_2_. Cells were passaged by 5 min incubation with TrypLE (Gibco) following wash with 1xPBS. Cells were passaged upon reaching approximately 80% confluence. Cells from passages 40-46 were used for the experiments. Prior to cell encapsulation, cells were detached from T75 flasks, centrifuged at 1200 rpm, resuspended in media and counted. Required number of cells per gel volume were then recentrifuged into pellets prior to resuspending in DexVS or HAVS SpyCatcher003 polymer precursor solutions.

### Substrate Coating and Blocking

Microplate wells were coated with 10 µg/mL of target protein solutions in 1x PBS or TBS storage buffer (Fn and Vn crosslinkers) and adsorbed to surface via physisorption. Proteins included full-length Fibronectin (Fn), ST-Fn-ST, ST-Fn 9,10 ΔRGD-ST, Vitronectin (Vn), Vn-L, Vn-S, and BSA controls and were incubated for 1 h at 37 °C in a humidified incubator. To eliminate non-specific cell adhesion, wells were blocked with 1% (w/v) heat-denatured bovine serum albumin (HD-BSA) in PBS for 30 min at 37 °C and subsequently washed twice with phosphate-buffered saline (PBS).

### Cell Adhesion Assay

Mouse Embryonic Fibroblasts (MEFs) were harvested and resuspended in serum-free medium. Cells were seeded at a density of 27,000 cells/cm² and incubated for 20 min at 37 °C in a humidified atmosphere containing 5% CO2. Non-adherent cells were gently removed by washing twice with PBS supplemented with 0.9 mM Ca²⁺ and 0.5 mM Mg²⁺ (PBS^++^).

### CyQUANT Fluorometric Assay

Parallel plates were stored at −80 °C to promote cell membrane lysis. Relative cell densities were quantified using the CyQUANT Cell Proliferation Assay Kit (ThermoFisher Scientific, Cat# C7026) according to the manufacturer’s protocol. Fluorescence intensity was measured at excitation and emission wavelengths of 480 nm and 520 nm, respectively, using a Tecan Spark multimode microplate reader. All experiments were performed in triplicate (n = 3).

### Crystal Violet Quantification

Adherent cells were fixed and stained using 0.5% (w/v) Crystal Violet in 10% methanol for 1 h at room temperature. Unbound dye was thoroughly rinsed with deionized water, and the plates were air-dried. Bound dye was extracted using 10% (v/v) acetic acid, and absorbance was measured at 570 nm on a microplate reader. n=4 replicates throughout assay.

### Microscopy and Image Processing

Brightfield morphological images of the 2D cell cultures were captured on an Olympus IX51 inverted microscope, and 3D matrix cultures on a Nikon Ti2 inverted fluorescent microscope. Fluorescent images of DAPI-stained nuclei were acquired using an Evident FV4000 laser scanning confocal microscope. Scale bars represent 200 µm for 2D culture brightfield images and DAPI-stained confocal images, and 50 µm for 3D matrix culture images.

### Live/Dead Staining

HCT116 cells were encapsulated in Dex and HA based matrices and cultured for 7 days in complete DMEM. On day 7, medium was gently removed and matrices were washed with 1xPBS. A solution of 2 µM calcein AM (Thermofisher Scientific) and 1 µM ethidium bromide (Thermofisher Scientific) in 1xPBS was added to the wells. Matrices were incubated for 20 min and then imaged with a Nikon Ts2 epifluorescence microscope at 10x magnification on 494 nm (Calcein AM live) channel and 528 nm (EtBr dead) wavelength. Post processing was conducted with Fiji ImageJ with addition of scale bars.

### Dome Invasion Model

Dex and HA hydrogel precursor solutions were prepared as previously described. Dome invasion models were created with droplets of a cell containing10 µL inner dome surrounded by blank 40 µL outer droplet domes. For polymer backbone studies cells were seeded at 1 million cells/mL density within inner dome (10,000 cells in 10 µL domes). Constructs were analyzed over culture period of two weeks with periodic brightfield imaging using Nikon Ts2 microscope.

### Spheroids in Viscoelastic Matrices

HCT116 cells were encapsulated in DexVS hydrogel precursor solutions supplemented with RatCol™ Rat Tail Collagen Type I (Advanced Biomatrix, REF: 5153) prepared as previously described, and plated onto 50 or 100 µg/ml poly-D-lysine (Gibco, REF: A3890401) coated Ibidi µ-Slide 15 Well 3D glass-bottom coverslips (REF: 81506). Images of the matrices were taken on days 1 and 7, with minimum three images and 10 spheroids per well imaged for analysis. Data was collected from n = 3 biological replicates, each with a minimum of n = 3 technical replicates for all groups. Image analysis was conducted in FIJI (ImageJ Version 1.54t), by manually thresholding each spheroid followed by particle analysis of all particles larger than 40 µm^2^ with a circularity between 0.30 -1.00.

### Statistical Analysis

All statistical analyses were carried out using GraphPad Prism version 10. Data are expressed as mean ± SD. Statistical significance was defined as p < 0.05 and is indicated in the figures by asterisks: * (p < 0.05), ** (p < 0.01), *** (p < 0.001), and **** (p < 0.0001). Group comparisons were assessed using one-way analysis of variance (ANOVA) with Tukey’s multiple-comparison post hoc test unless otherwise specified. For cell adhesion assays, data distribution normality was evaluated using the Shapiro-Wilk test. For non-normally distributed crystal violet datasets (N = 4), statistical differences among groups were evaluated using a non-parametric Kruskal-Wallis test followed by Dunn’s post-hoc test. For normally distributed CyQUANT datasets (N = 3), a one-way Analysis of Variance (ANOVA) followed by Bonferroni’s post-hoc test was performed. The specific statistical approaches used for each experiment are described in the associated figure legends. All presented images and graphical data are representative of repeated experiments, as detailed in the corresponding methods sections and figure legends.

## Resource Availability

All data and materials are accessible from corresponding author upon reasonable request.

## Acknowledgements

The authors gratefully acknowledge Biocenter Finland for providing research infrastructure support. We also thank Niklas Kähkönen and Janne Kärnä (Tampere University) for their valuable technical assistance. This work was supported by the Sigrid Juselius Foundation (project nos. 240150 to O.P.O. and 230140 to V.P.H.), the Finnish Cancer Foundation (Syöpäsäätiö), the Linnea Pirilä Fund and Aino Valvaalan Fund of the Finnish Cultural Foundation Pirkanmaa Regional Fund (grants 50241473 and 50251179 to A.D.E.), Tampere Tuberculosis Foundation (P.T.), the Tampere University Doctoral School Fund (A.D.E.), the Cancer Society of Finland (V.P.H.), and the Research Council of Finland (grants 363941 and 331946 to V.P.H.; 339449 and 363616 to R.R.). J.K. is a doctoral researcher in the iCANDOC Doctoral Education Pilot in Precision Cancer Medicine. Figures were prepared using Biorender.com and PerkinElmer ChemDraw Professional.

## Author Contributions

Conceptualization, A.D.E., P.T., R.R., O.P.O., and V.P.H.; methodology, A.D.E., J.K., S.R., M.M., H.H., P.T., R.R., O.P.O. and V.P.H.; validation, A.D.E., J.K., S.R., M.M., and H.H.; formal analysis, A.D.E., J.K., S.R., M.M., and H.H.; investigation, A.D.E., J.K., S.R., M.M., and H.H.; resources, P.T., R.R., O.P.O., and V.P.H.; data curation, A.D.E., J.K., S.R., M.M., and H.H.; writing – original draft, A.D.E., J.K., R.R. and V.P.H.; writing – review & editing, A.D.E., J.K., S.R., M.M., H.H., P.T., R.R., O.P.O, and V.P.H., and; visualization, A.D.E., J.K., S.R., M.M., and H.H.; supervision, P.T., O.P.O., R.R., and V.P.H.; project administration, O.P.O., R.R., and V.P.H.; funding acquisition, A.D.E., J.K., P.T., R.R., O.P.O, and V.P.H.

## Declaration of Interests

A.D.E, R.R., P.T, S.R, O.P.O, and V.H. have filed a PCT patent application (PCT/FI2026/050362) related to the work reported in this manuscript. All other authors declare no competing interests.

