## Supplementary Information for "Genetically programmable molecular framework for constructing precision recombinant human ECM mimics"

### Supplementary Materials

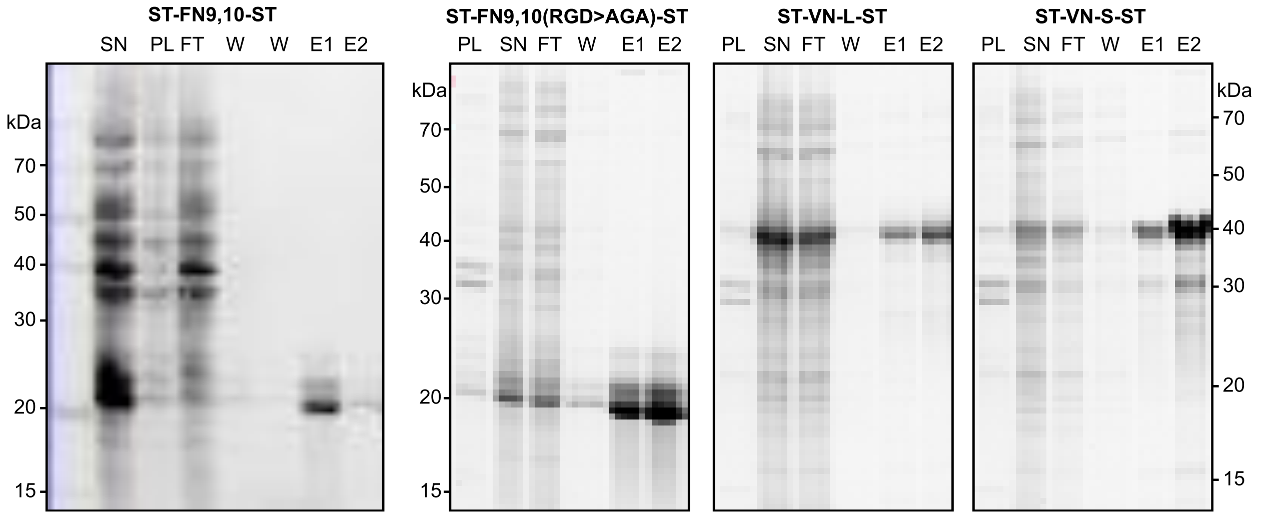


**Supplementary Figure S1: Recombinant protein ECM crosslinkers.** SDS-PAGE gels after expression and purification of SpyTag-fibronectin-9,10-SpyTag, SpyTag-fibronectin-9,10(ΔRGD>AGA)-SpyTag, SpyTag-vitronectin-L-SpyTag, and SpyTag-vitronectin-S-SpyTag (from left to right) The three gels on the right were run on the same SDS-PAGE gel with image cropping to separate individual adapters to improve clarity. SN, soluble fraction, PL, insoluble fraction, FT, unbound fraction, W, wash sample, E, elution fractions.

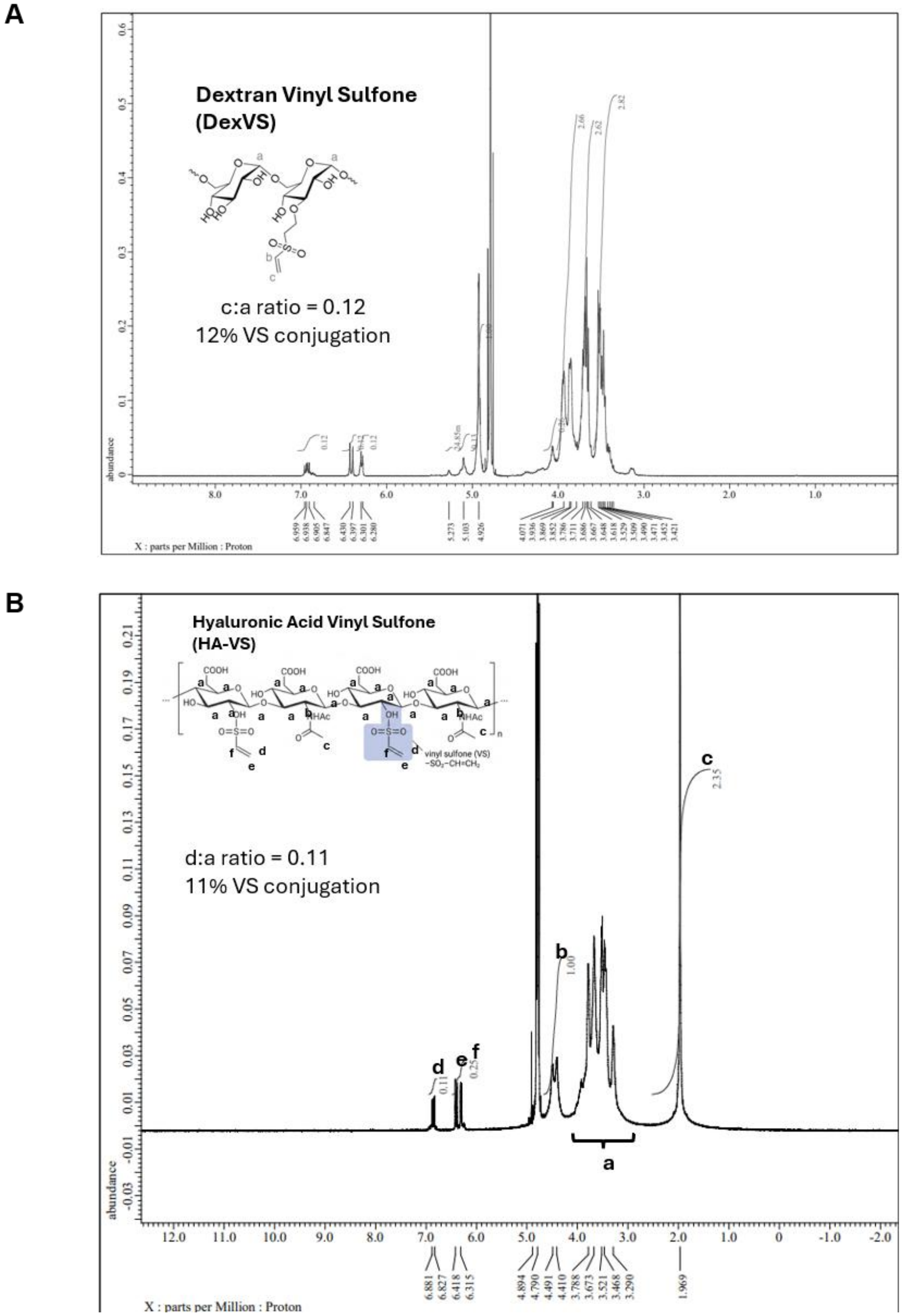


**Supplementary Figure S2: Vinyl sulfone conjugation A.** $^{1}$H-NMR (500 MHz) spectra of dextran vinyl sulfone recorded in D2O at 298K. **B.** $^{1}$H-NMR (500 MHz) spectra of Hyalronic acid vinyl sulfone recorded in D2O at 298K.

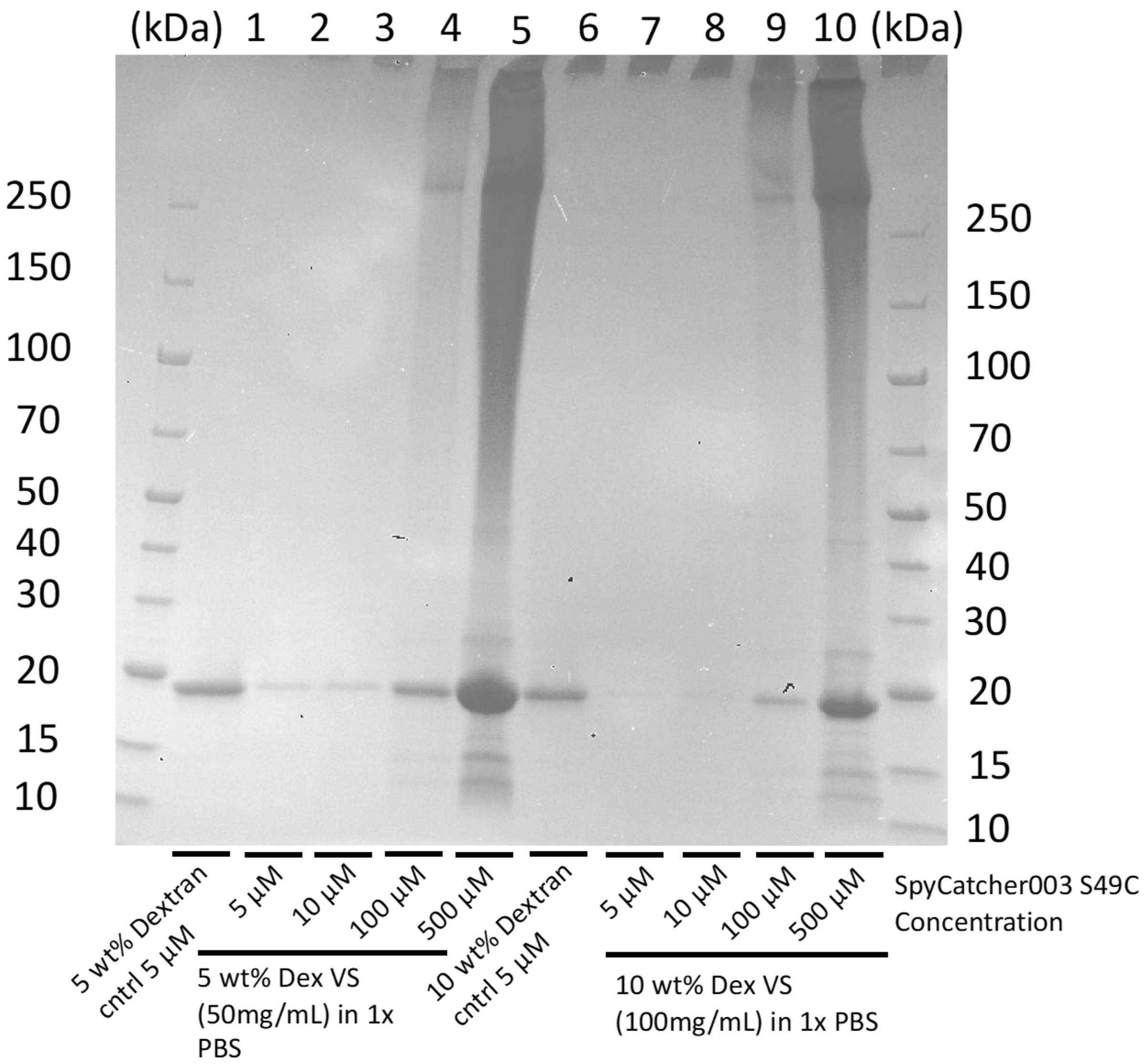


**Supplementary Figure S3: Binding of Spycatcher003 S49C to DexVS backbone** with 1 hour incubation (left) and 24-hour incubation (right) with different concentrations of DexVS and Spycatcher003 S49C using Coomassie Blue SDS-PAGE gels.

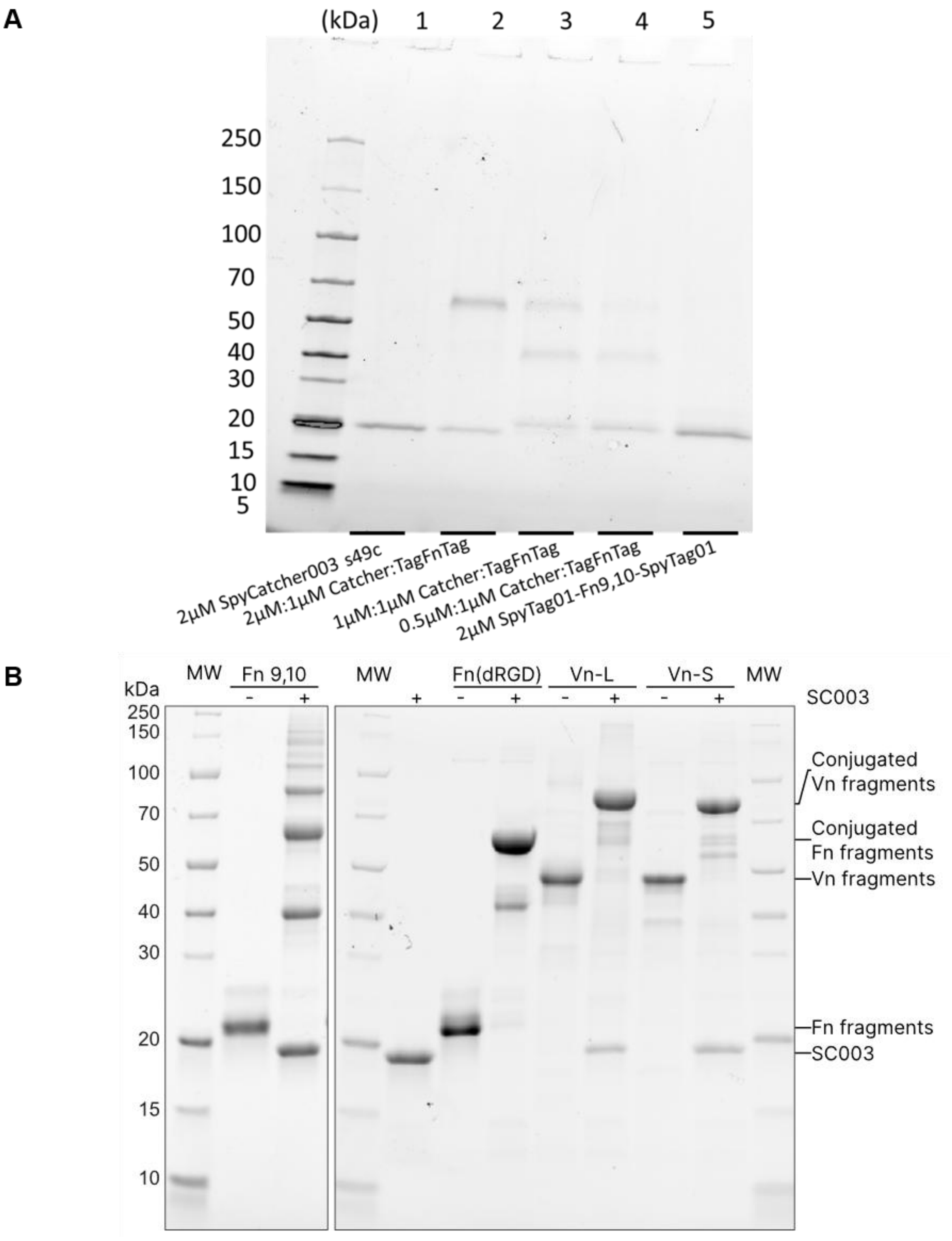


**Supplementary Figure S4: Demonstration of isopeptide bond formation between SpyCatcher and pytagged ECM crosslinkers. A.** Stain-Free SDS-PAGE band shift assay of SpyCatcher003 S49C conjugation to SpyTag01-Fn9,10-Spytag01 at varying ratios. **B.** Band shift assay for SpyTag mediated conjugation of ECM crosslinker- SpyTag-fibronectin-9,10-SpyTag, SpyTag-fibronectin-9,10(ΔRGD>AGA)-SpyTag, SpyTag-vitronectin-L-SpyTag, and SpyTag-vitronectin-S-SpyTag to SpyCatcher003 S49C. The later three were part of same SDS-PAGE gel.

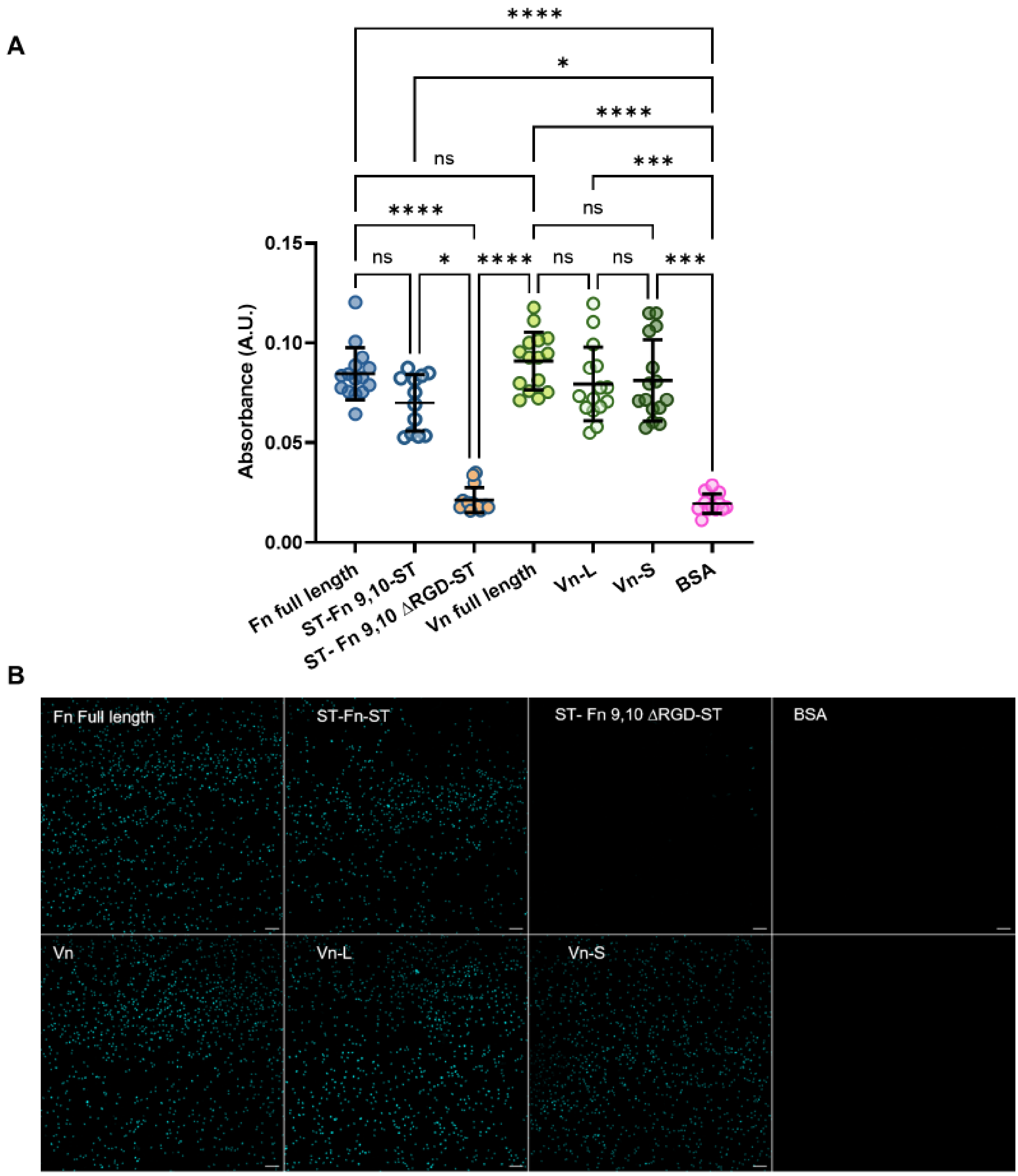


**Supplementary Figure S5: Crystal violet and CyQUANT visualization cell adhesion extended data A.** Relative cell adhesion quantified by Crystal Violet dye extraction (Absorbance at 570 nm; n = 4 independent biological replicates). Data is presented as mean ± SD. Data was analyzed using Kruskal-Wallis test with Dunn's post-hoc test due to non-normal distribution. Statistical significance was defined as $p < 0.05$ and is indicated in the figures by asterisks: * ($p < 0.05$), ** ($p < 0.01$), *** ($p < 0.001$), and **** ($p < 0.0001$). **B.** Representative images of cell adhesion to ECM adapters following CyQuant staining, scale bar = 200 µm.
